# Striatal direct and indirect pathway neurons differentially control the encoding and updating of goal-directed learning

**DOI:** 10.1101/2020.02.18.955385

**Authors:** James Peak, Billy Chieng, Genevra Hart, Bernard W. Balleine

**Affiliations:** Decision Neuroscience Lab, School of Psychology, UNSW Sydney, Australia

**Keywords:** Instrumental conditioning, goal-directed learning, goal-directed action, posterior dorsomedial striatum, direct pathway neurons, indirect pathway neurons, spiny projection neurons

## Abstract

The posterior dorsomedial striatum (pDMS) is necessary for goal-directed action, however the role of the direct (dSPN) and indirect (iSPN) spiny projection neurons in the pDMS in such action remains unclear. In this series of experiments, we examined the role of pDMS SPNs in goal-directed action and found that, whereas dSPNs were critical for goal-directed learning and for energizing the learned response, iSPNs were involved in updating that learning to support response flexibility. Instrumental training elevated expression of the plasticity marker Zif268 in dSPNs only, and chemogenetic suppression of dSPN activity during training prevented goal-directed learning. Unilateral optogenetic inhibition of dSPNs induced an ipsilateral response bias in goal-directed action performance. In contrast, although initial goal-directed learning was unaffected by iSPN manipulations, optogenetic inhibition of iSPNs, but not dSPNs, impaired the updating of this learning and attenuated response flexibility after changes in the action-outcome contingency.

## Introduction

Animals adapt to changing environments and maximize opportunities for reward by flexibly adjusting their actions according to current goals. Such goal-directed actions have been found to rely on the ability of animals to encode the relationship between an action and its consequences or outcome, and to integrate that information with the current value of the outcome (Balleine & Dickinson, 1998; Dickinson & Balleine, 1994). It has also become clear that the acquisition of goal-directed actions relies on a corticostriatal circuit centered on the posterior region of dorsomedial striatum (pDMS) where glutamatergic inputs from the prelimbic cortex and dopaminergic inputs from the substantia nigra pars compacta (SNc) interact to influence goal-directed learning and performance via their effect on striatal output (Shiflett & Balleine, 2010; Peak *et al*, 2019; Balleine, 2019, for reviews). Accordingly, in the past decade there has been intense investigation into how striatal output pathways mediate the expression of actions, goal-directed or otherwise (Poyraz *et al.*, 2016; Freeze *et al.*, 2013; Hikida *et al.*, 2010; Kravitz *et al.*, 2010; Kravitz *et al.*, 2012; Tai *et al.*, 2012; Tecuapetla *et al.*, 2014; Tecuapetla *et al.*, 2016). Nevertheless, despite these considerable efforts, consensus on this issue has proven elusive.

The analysis of striatal output has long focused on the influence of two pathways, one projecting directly to substantial nigra pars reticulata (SNr) and another indirectly to the SNr via the external globus pallidus (GPe) and subthalamic nucleus (STN) (Albin *et al.*, 1989). Canonically, it has been argued that each pathway is controlled by a distinct population of spiny projection neuron; the direct pathway SPNs (dSPNs), that predominantly express the dopamine D1 receptor (D1R), and the indirect pathway SPNs (iSPNs), that express the dopamine D2 receptor (D2R) (Gerfen & Surmeier., 2011; Calabresi *et al.*, 2014 for reviews). As a consequence, experiments investigating the distinct functions of striatal output pathways have typically relied on the effects of recording, imaging or manipulating the activity of these distinct cell types as proxies for the output pathways themselves. These studies have been inconclusive. Within the pDMS, goal-directed learning has been found to induce plasticity at dSPNs (Shan *et al.*, 2014, Maroteaux *et al.*, 2014; Matamales *et al.*, 2020) which also appear to control goal-directed performance; unilateral stimulation of dSPNs in the DMS drives a contralateral response bias (Tai *et al.*, 2012). By contrast, the function of D2-expressing SPNs has proven ambiguous; whereas most studies suggest little involvement in goal-directed learning (Sippy *et al.*, 2015), their role in performance has ranged variously from no influence, to a cooperative influence (Cui *et al.*, 2013; Tecuapetla *et al.*, 2016), to an inhibitory influence on action selection and vigor (Poyraz *et al.*, 2016; Tai *et al.*, 2012).

More recently, evidence has emerged that questions the use of a cell-type specific approach to investigate the functions of these output pathways. Although there has been a thick veneer of agreement in the literature with the canonical view, considerable evidence has accumulated questioning the exclusivity of the influence of D1- and D2-expressing SPNs on the direct and indirect output pathways, respectively. Over and above evidence for coexpression of D1 and D2 receptors on SPNs (Bertran-Gonzalez *et al.*, 2008, Gagnon *et al.*, 2017), evidence from single neuron tracing studies suggests considerable colateralisation, with D1 expressing neurons projecting to SNr but also to GPe (Cazorla *et al.*, 2014; Wu, Richard & Parent, 2000; Fujiyama *et al.*, 2011; Kawaguchi *et al.*, 1990; see Burke *et al.*, 2017 for review). Furthermore, although a number of studies have previously documented modest D2-to-D1 neuron interactions within the striatum (Taverna *et al.*, 2008), recent evidence suggests that the convergence between striatal systems has been underestimated and is far more prevalent than previously thought (Matamales *et al*, 2020), providing a form of intrastriatal transmodulation capable of shaping the functional role of large striatal territories. The general conclusion from these studies is, therefore, that, given the degree of interaction between D1- and D2-expressing SPNs, the function of striatal output pathways may be more accurately evaluated based on the target of projection neurons rather than their cell type.

The current series of experiments sought, therefore, to evaluate the functions of the specific populations of projection neurons in the striatum based on their targets, using the retrograde transport of Cre to target striatal neurons projecting to the SNr compared to those projecting to the GPe. We did not seek to segregate populations of projection neurons but to evaluate the functions of these projections in an unbiased manner on the basis of their projection target, whatever the degree of overlap. Using this approach, we report clear evidence of dSPN but not iSPN involvement in goal-directed learning and performance and of iSPN but not dSPN involvement in updating goal-directed learning when the action-outcome contingencies controlling goal-directed performance change.

## Results

### The acquisition of goal-directed action induces Zif268 expression in dSPNs in the pDMS

We first sought to establish whether plasticity-related activity differs in dSPNs and iSPNs during the acquisition of goal-directed action. Based on the above considerations, we developed a pathway-specific approach, targeting dSPNs and iSPNs by injecting the retrograde tracers, fluorogold (FG) and cholera-toxin B subunit (CTB), bilaterally into the direct monosynaptic targets of each pathway, the SNr and GPe, respectively (Figure 1A-B). We were able to identify dSPNs and iSPNs in the same animal; Figure 1C shows triple labeling of SPNs with either FG, CTB or both, co-labeled with DARPP-32, a marker for SPNs. Figure 1D shows retrograde tracer counts; there was no difference in number of cells/mm^2^ labelled with FG or CTB (F<1) or in the mean percentage of neurons co-labeled with either FG or CTB (F<1.7), suggesting that this approach was not significantly biased towards one or other population of SPNs. Overall, we were able to target approximately half of all SPNs identified with DARPP-32 (Figure 1E) although, confirming previous reports, there was substantial overlap of dSPN and iSPN populations (estimated at 16-17% of the total number of retrogradely labeled cells).

**Figure 1.**
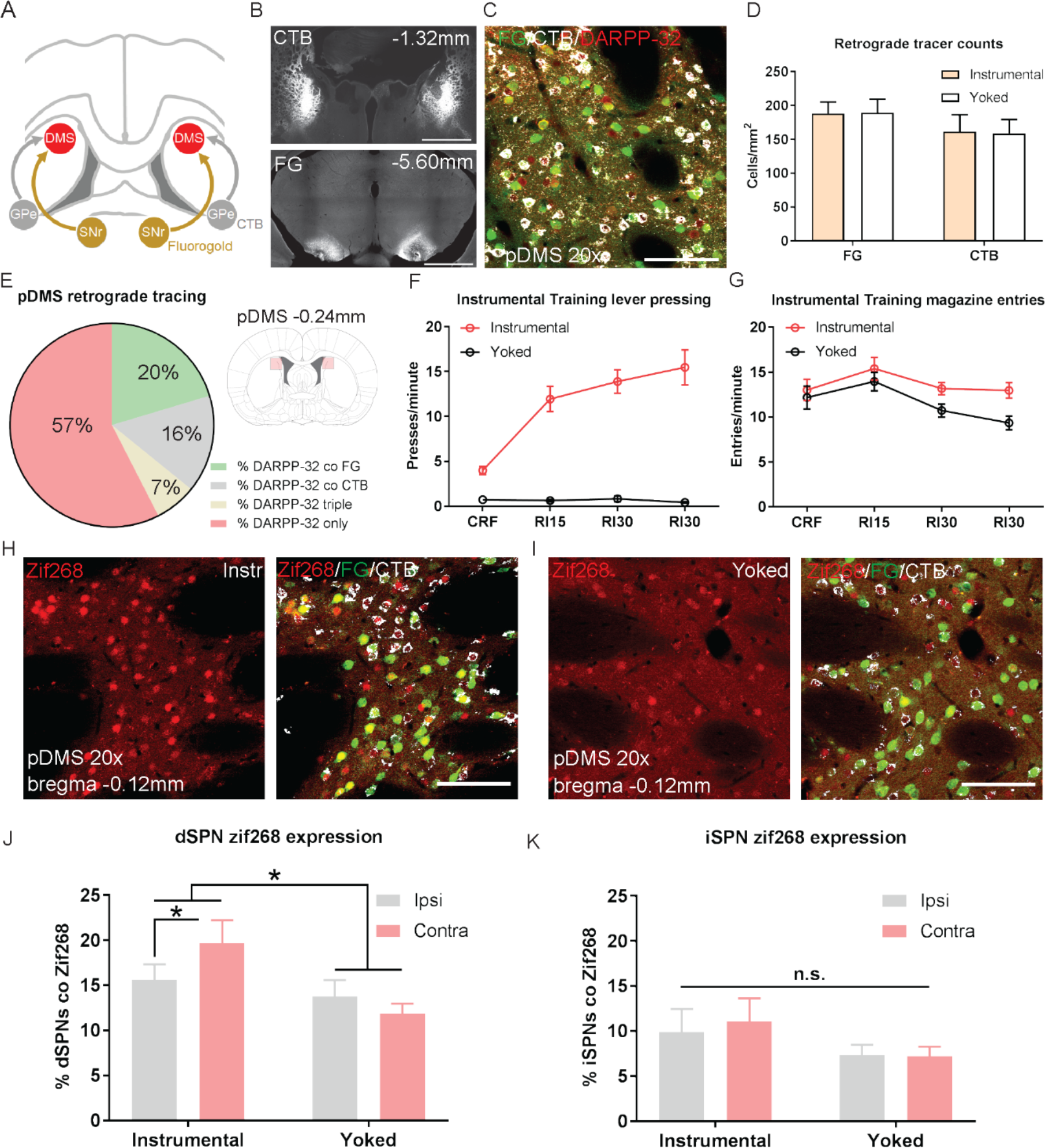
The acquisition of goal-directed actions induces response-specific expression of Zif268 in dSPNs in the pDMS. **A**, Schematic illustrating the surgery design for retrograde tracing. Rats received bilateral infusions of the retrograde tracers Fluorogold (FG) and Cholera-Toxin B (CTB) into the SNr and GPe respectively. **B,** Fluorescent confocal images showing injection sites of CTB (top) and FG (bottom) in the GPe and SNr respectively, scale bars, 2000μm. **C,** Fluorescent confocal image taken from one 40μm coronal section in the pDMS, illustrating fluorescence labelling of DARPP-32 (red), FG (green), CTB (white), scale bar, 100μm. **D,** Mean (± SEM) total number of cells labelled with FG and CTB for rats in Group Instrumental and Group Yoked. **E,** Percentages represent the proportion of total DARPP-32 positive SPNs that also express FG or CTB in the pDMS (imaging region indicated in pale red, right). Data presented are mean percentages for 23 rats, averaged across hemispheres. **F,** Mean (± SEM) lever presses per minute averaged across each day of instrumental or yoked training for each group. **G,** Mean (± SEM) magazine entries per minute averaged across each day of instrumental or yoked training for each group. **H,** Fluorescent confocal image of a coronal 40μm section for an instrumentally trained rat; left image shows Zif268 expression (red), and right image shows the same merged with FG labelled dSPNs (green) and CTB labelled iSPNs (white), scale bar, 100μm. **I,** Same as G but for a yoked trained rat, scale bar, 100μm. **J,** Percentage of labelled dSPNs that were co-labelled with Zif268 for rats in Group Instrumental and Group Yoked in the hemisphere ipsilateral to lever position (grey bars) and contralateral to lever position (pink bars). Data presented are group means of four pDMS sections per rat. **K,** Same as J but for iSPNs. All data presented are means ± SEM. \**p* < 0.05.

Next, we assessed the learning related activity in dSPNs and iSPNs induced by instrumental training. Half of the rats were given training over 4 days, in which pressing a lever (located to the left or right side of a central food magazine) delivered grain pellets on increasing random interval schedules of reinforcement (see methods). A second, control group, received yoked training in which grain pellets were delivered at the same average interval as the instrumentally trained rats, but independently of lever pressing. This resulted in robust acquisition of the instrumental response for rats in Group Instrumental, but not for rats in Group Yoked (Figure 1F; F(1,20)=90.728, *p*<0.001). Both groups entered the magazine to retrieve the food, and although group Instrumental had higher entry rates, this wasn’t significant (Figure 1G; F(1,20)=4.035, *p*=0.058).

Ninety minutes after the beginning of the final training session, rats were perfused and we used immunofluorescence to measure the expression of the immediate-early gene Zif268, widely used as an activity marker for learning-related plasticity (Davis *et al.*, 2000), in dSPNs and iSPNs in the pDMS. There was robust labeling of Zif268 in both groups (Figure 1H-I). We quantified Zif268 expression in each hemisphere separately, and grouped hemispheres according to their relative location to the lever in the chamber; i.e., if a rat had the lever on the right side of the chamber, the left hemisphere was designated as ‘contralateral’, and the right hemisphere as ‘ipsilateral’ (and vice versa).

Figures 1J and 1K show the percentage of dSPNs and iSPNs co-labelled with Zif268, respectively. Rats in Group Instrumental had significantly higher Zif268 expression in dSPNs than rats in Group Yoked (Figure 1J; F(1,20)=4.521, *p*=0.046). Furthermore, there was a pronounced hemispheric difference in Group Instrumental but not in Group Yoked; rats in Group Instrumental had significantly more Zif268 in the hemisphere contralateral to the lever side, indicating that this difference was specific to instrumental training (group x hemisphere interaction, (F(1,20)=4.836, *p*=0.040). In contrast, analysis of Zif268 expression in iSPNs (Figure 1K) revealed no significant differences between Group Instrumental and Group Yoked in overall levels of Zif268 expression, and no differences between hemispheres (Fs<1.5).

### Goal-directed learning is disrupted by chemogenetic inhibition of dSPNs but not iSPNs in the pDMS

Having established that instrumental training differentially alters the intracellular activity of dSPNs, we next sought to assess the causal involvement of dSPNs and iSPNs in goal-directed learning. In order to induce prolonged suppression of neuronal activity across each day of instrumental training, chemogenetic (DREADDs) inhibition is optimal; however, the use of DREADDs in striatal neurons has been a topic of debate in the literature with concerns raised over their effectiveness. Although there have been successful reports of the use of DREADDs in striatal neurons (Ferguson *et al.*, 2011; Ferguson *et al.*, 2013), SPNs have a relatively low number of G-protein-activated inwardly rectifying potassium (GIRK) channels (Lovinger, 2010). We therefore assessed whether hM4Di DREADDs expressed on SPNs could suppress cortically-evoked neuronal firing following the application of the synthetic ligand of the hM4Di receptor, clozapine-n-oxide (CNO).

We used a dual-virus approach (Figure 2A; Marchant *et al.*, 2015; Hart *et al.*, 2018a) to express hM4Di receptors on dSPNs or iSPNs, by bilaterally infusing a retrogradely transported AAV-virus expressing Cre recombinase (rAAV5.CMV.HI.eGFP-CRE.WPRE.SV40; retro-Cre) into either the SNr or GPe, so as to express Cre on dSPNs or iSPNs, respectively. We then injected a Cre-dependent hM4Di DREADDs virus (rAAV5/hSyn-DIO-hM4D(Gi)-mCherry; DIO-hM4D) into the pDMS, to selectively express DIO-hM4D on either dSPNs or iSPNs. To induce neuronal firing in SPNs, we injected channelrhodopsin (AAV5-CaMKIIa-hChR2(H134R)-eYFP; ChR2-eYFP) into the prelimbic cortex (Figure 2A), which has dense bilateral glutamatergic projections onto dSPNs and iSPNs (Hart *et al.*, 2018a; Wall *et al.*, 2013). We used slice electrophysiology to assess the effect of CNO on the cortically-evoked firing of dSPNs and iSPNs expressing hM4Di DREADDs.

**Figure 2.**
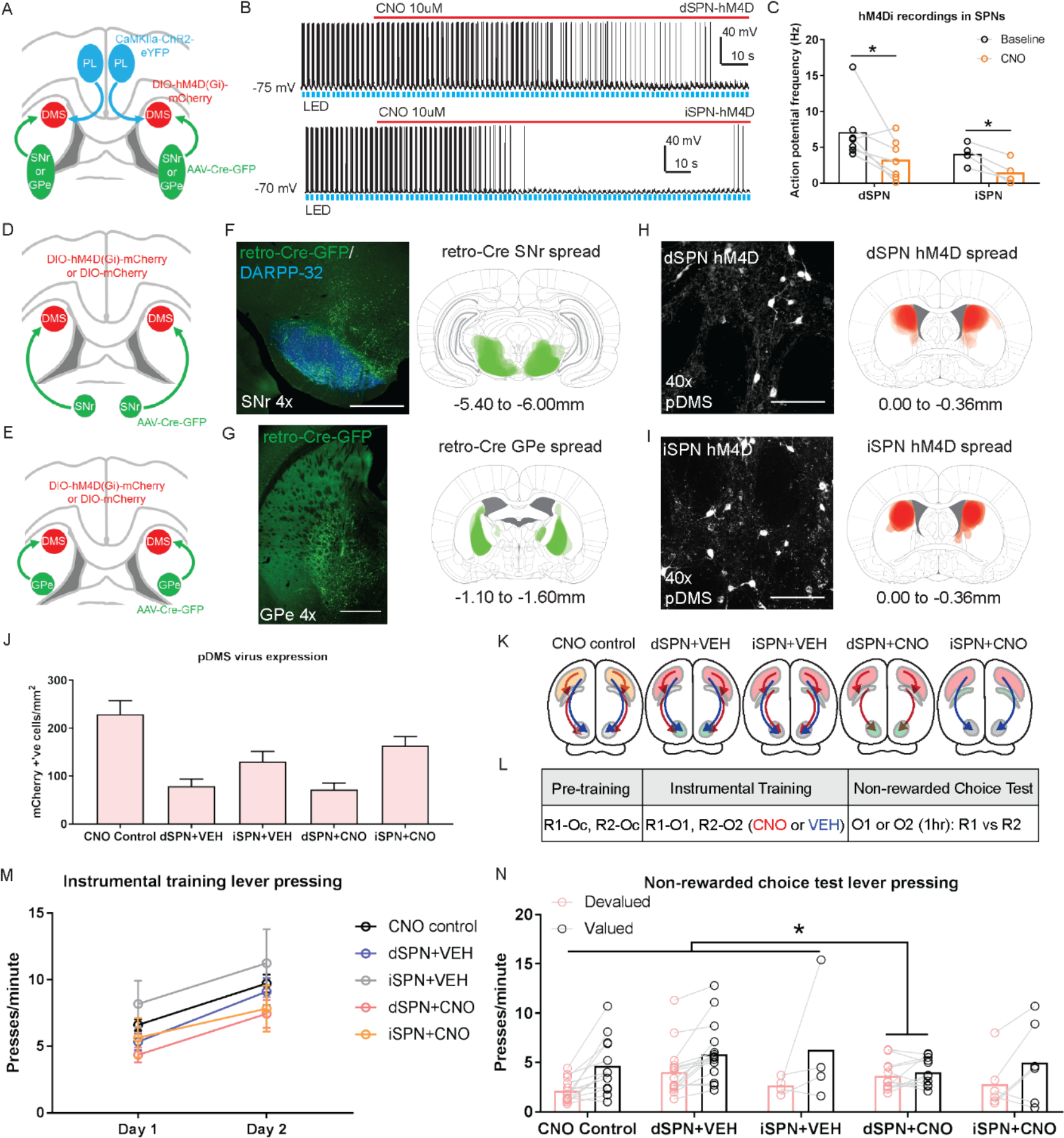
Goal-directed learning is disrupted by chemogenetic inhibition of dSPNs but not iSPNs in the pDMS. **A,** Schematic depicting the viral injections for electrophysiology recordings; retro-Cre was injected into the SNr or GPe, DIO-hM4D was injected into the pDMS and ChR2-eYFP injected into the PL; direction of arrows indicates retrograde or anterograde transport of the virus. **B,** Example trace from one recorded dSPN (top) and one recorded iSPN (bottom) expressing DIO-hM4D. Upward deflection black lines represent action potentials, which were elicited by the pulsing of a 473nm wavelength LED light (blue) onto corticostriatal terminals (see Methods). The red line indicates the time period when 10uM CNO solution was added to the extracellular solution. **C,** Action potential frequency of dSPNs and iSPNs elicited by light-evoked terminal glutamate release from ChR2-containing corticostriatal axons in the pDMS under baseline conditions (firing rate before CNO application) and after the application of CNO in the extracellular solution. Data presented are means from all recorded cells ± SEM. **D,** Schematic depicting the viral injections to target dSPNs; retro-Cre was injected bilaterally into the SNr and DIO-hM4D or DIO-mCherry was injected bilaterally into the pDMS. **E,** Same as D but for targeting iSPNs; retro-Cre was injected into the GPe. **F,** Confocal image (scale bar, 1000
μm) from one rat showing retro-Cre expression in the SNr (left) and the extent of virus spread in the SNr, imaged at the widest point for all included animals (right). **G,** Same as C but showing retro-Cre expression in the GPe. **H,** Confocal image (scale bar, 100μm) from one rat showing DIO-hM4D expression in pDMS dSPNs (left) and the extent of DIO-hM4D spread in the DMS (right). **I,** Same as H but showing DIO-hM4D expression in pDMS iSPNs. **J,** Total number of mCherry positive cells/mm^2^ in the pDMS in rats from each experimental group. **K,** Summary of experimental groups; Blue arrows indicate intact direct pathway function and red arrows indicate intact indirect pathway function. **L,** Summary of the experimental design; R1 and R2 indicate left and right lever responses; Oc, O1 and O2 indicate distinct food outcomes; CNO and VEH indicate injections of clozapine-N-oxide or vehicle, respectively. **M,** Mean (± SEM) lever presses per minute averaged across each day of instrumental training for each group. **N,** Mean lever presses per minute on the devalued and valued lever for each rat in each group, averaged across two days of a non-rewarded choice tests, \**p* < 0.05.

Figure 2B shows the raw trace recorded from a dSPN (top) and an iSPN (bottom) expressing DIO-hM4D; action potentials were elicited by pulsing an LED light onto ChR2-containing corticostriatal axons (blue bars). Following the administration of CNO, action potential frequency was significantly reduced (Figure 2C; dSPNs, t(6)=2.662, *p*=0.037; iSPNs, t(3)=4.921, *p*=0.016). We used the same dual-virus approach to express either DIO-hM4D or a control fluorophore lacking the hM4D construct (rAAV5/hSyn-DIO-mCherry; DIO-mCherry) on dSPNs (Figure 2D) or iSPNs (Figure 2E). Retro-Cre expression in the SNr and GPe are shown in Figure 2F and 2G, respectively. Figure 2H shows an example image of DIO-hM4D expression on pDMS dSPNs, and the spread of DIO-hM4D expression in all included animals (right). The same measures are shown for iSPNs in Figure 2I.

We quantified the mean number of cells expressing DIO-hM4D or DIO-mCherry for each animal (Figure 2J); there was no difference in the number dSPNs or iSPNs expressing DIO-hM4D (dSPN+VEH vs dSPN+CNO, F<1; iSPN+VEH vs iSPN+CNO, F<1). There were, however, significantly more SPNs (half dSPNs and half iSPNs) expressing DIO-mCherry (Group CNO control vs rest; F(1,46)=23.930, *p*<0.001). We compared the rate of infection achieved with this dual-virus approach in dSPNs to that obtained using retrograde tracing (Figure 1D); retro-Cre in combination with DIO-hM4D achieved 38-42% of the labelling achieved with fluorogold, and this percentage is consistent with that observed using the same viral combination in the PL-pDMS pathway, and compared to fluorogold (30-45% estimated by Hart *et al.*, 2018a). For iSPNs, retro-Cre in combination with DIO-hM4D achieved 81-102% of the labelling that was achieved with CTB. The substantially higher infection rate from GPe is likely to be due to a combination of two things; fewer CTB labelled neurons compared to FG (see Figure 1D) and more efficient retro-Cre transport in striatopallidal neurons, possibly due to the shorter pathway length.

For behavioral testing there were five groups of rats, summarized in Figure 2K. There were three control groups: (1) Group CNO control, expressed DIO-mCherry on SPNs (half on dSPNs and half on iSPNs which were pooled based on a lack of significant differences in press rate during training or test; Fs<3.7); (2) Group dSPN+VEH and (3) Group iSPN+VEH both of which expressed DIO-hM4D on dSPNs or iSPNs, respectively, and received injections of vehicle throughout training. The two experimental groups expressed DIO-hM4D and received injections of CNO throughout training to suppress the activity of dSPNs (Group dSPN+CNO) or iSPNs (Group iSPN+CNO).

A summary of the behavioural design is presented in Figure 2L. Rats were given 4 days of instrumental pre-training with two levers for a common outcome (Oc) after which they received two days of instrumental training with the same two levers paired with new, distinct outcomes (R1-O1, R2-O2). Intraperitoneal injections of CNO or VEH were administered one hour prior to these training sessions. All rats acquired the instrumental response across pre-training (Figure S1A). Across two days of instrumental training (Figure 2M), all rats increased their response rates from Day 1 to Day 2 (F(1,46)=83.276, *p*<0.001) and there were no significant differences between groups in overall response rates (Bonferroni, k=3, highest F=4.856, dSPN vs controls).

In order to assess goal-directed learning, we gave rats a choice extinction test, drug free, immediately after specific satiety-induced outcome devaluation. For this devaluation, rats were allowed free access to one of the previously earned outcomes (O1 or O2) for one hour after which they were returned to the operant chambers and tested with both levers available in a choice extinction test. Goal-directed action is demonstrated by rats showing a reduction in responding on the lever associated with the now-devalued outcome relative to the other lever, indicating their ability to integrate prior action-outcome learning with the current value of the outcome. Responding on the valued and devalued levers is presented for each rat in each group in Figure 2N. We compared responding in Groups dSPN+CNO or iSPN+CNO against the three control groups; dSPN+VEH, iSPN+VEH and CNO control. We also compared the control groups that received vehicle to the CNO control. There were no differences between groups in overall rates of responding (main effect of group, Fs<2.1). There was, however, a main effect of devaluation (Bonferroni, k=3, F(1,46)=28.726, *p*<0.001) and, importantly, a significant group (dSPN+CNO vs controls) x devaluation interaction (Bonferroni, k=3, F(1,46)=6.930, *p*=0.012) suggesting that, whereas the control groups showed a reliable devaluation effect, this was abolished in rats that had dSPNs suppressed during instrumental training. Importantly, there was no difference in the magnitude of the devaluation effect in Group iSPN+CNO relative to the three control groups, or between the two vehicle groups and Group CNO control (Fs<1). These results suggest that specifically dSPN activity in the pDMS is necessary during training for animals to acquire goal-directed instrumental actions.

In a subset of rats, we examined whether this effect was attributable to a diminished capacity to discriminate between responses or outcomes, when dSPNs or iSPNs were suppressed. To achieve this, rats were injected with CNO or vehicle (group allocations were unchanged) and given a rewarded choice test following outcome devaluation (Figure S1B), in which each lever delivered its respective outcome, negating the need for animals to use prior learning to bias actions. Under these circumstances, all rats were able to respond according to current outcome value, and there were no differences between groups (main effect of lever, Bonferroni, k=3, F(1,18)=53.237, *p*<0.001; no effect of group and no group x lever interactions, Fs<1.6). This result also demonstrates that specific satiety-induced outcome devaluation is effective under dSPN (or iSPN) inhibition. Together, therefore, these results suggest that dSPNs are selectively involved in encoding the specific action-outcome associations that support goal-directed learning during instrumental training.

Finally, although we saw no evidence for motor disruption across training or during the rewarded choice test when activity in dSPNs or iSPNs was suppressed, we assessed whether suppressing activity in pDMS SPNs induces a motoric impairment more directly using a rotarod test (Hamm *et al.*, 1994). Rats infused with DIO-hM4D were all tested twice; once under vehicle and once under CNO. As shown in Figure S1C, suppressing SPNs did not affect the amount of time spent on the rotarod for rats in any group (pairwise comparisons between CNO and VEH in each group, highest F=2.7, dSPN+VEH). Therefore, the suppression of dSPNs or iSPNs produced no overt motoric impairments that were detectable using this test.

### Chemogenetic stimulation of iSPNs in the pDMS leaves goal-directed learning intact

Although the previous results suggest goal-directed learning relies on dSPN plasticity during training, it is possible learning was disrupted by altering the relative output of dSPNs and iSPNs to favor heightened iSPN output. Therefore, we tested whether increased iSPN activity could similarly affect goal-directed learning. We used an hM3Dq DREADD to stimulate the activity of iSPNs and slice electrophysiology to verify that CNO increased neuronal firing in iSPNs expressing Cre-dependent hM3Dq DREADDs (rAAV5/hSyn-DIO-hM3D-mCherry; DIO-hM3D), targeted using the same retro-Cre approach described (Figure 3A-B). This was confirmed; application of CNO produced a significant increase in the number of action potentials recorded from iSPNs expressing DIO-hM3D, relative to baseline (Figure 3C-D; paired t-test, \**p*<0.05).

**Figure 3.**
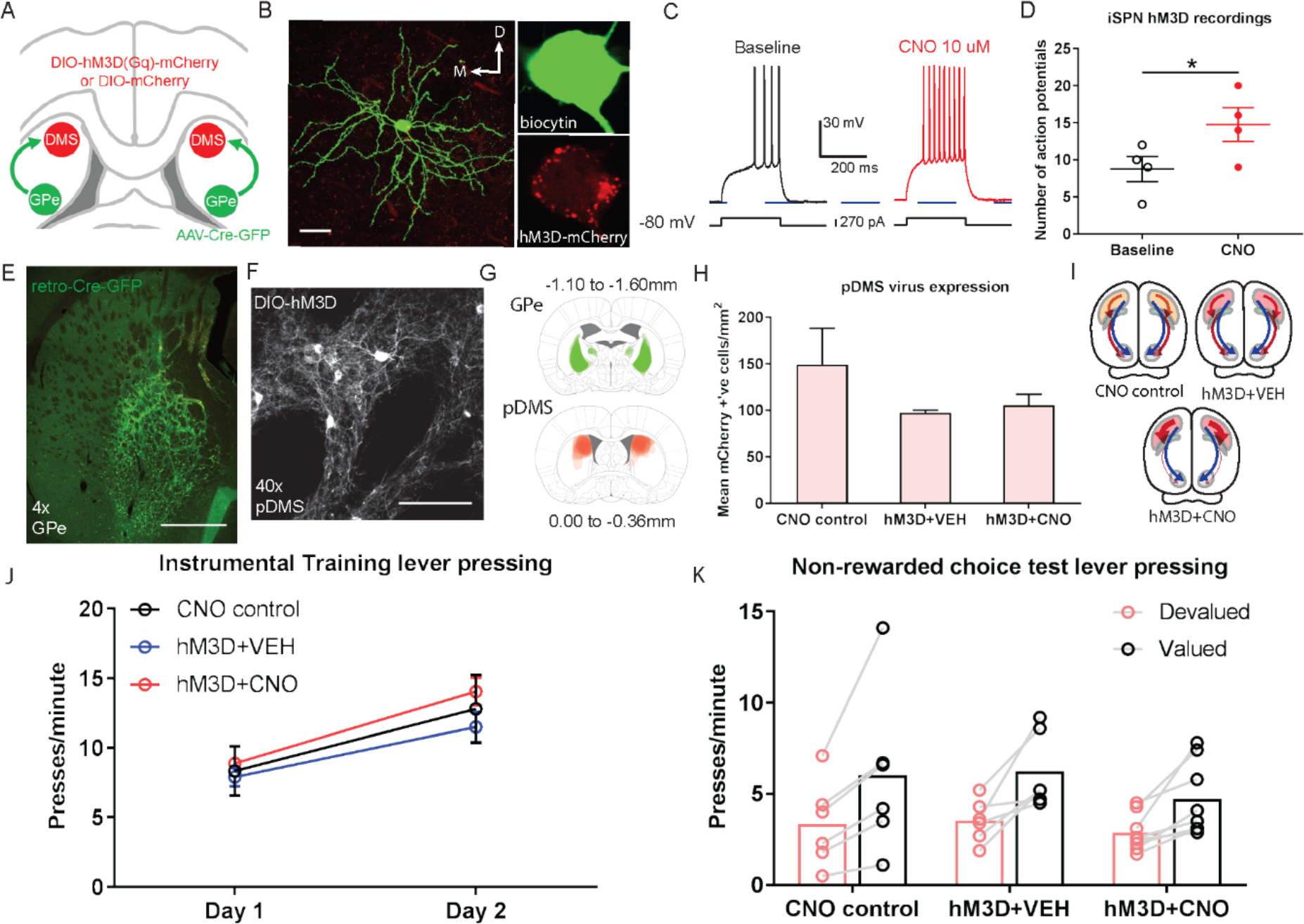
Chemogenetic stimulation of iSPNs in the pDMS leaves goal-directed learning intact. **A,** Schematic depicting the viral injections, retro-Cre was injected bilaterally into the GPe and DIO-hM3D or DIO-mCherry was injected bilaterally into the pDMS. **B**, An example biocytin-labelled iSPN expressing hM3D from recording (scale bar, 30μm). **C,** An example trace from one iSPN (pictured in B) from injection of a depolarizing current step (at resting membrane potential) before and after bath application of CNO. **D,** Number of action potentials evoked from an identical size current step injection before and during CNO application in hM3D-expressing iSPNs. The chosen step size in each neuron was when current injection first elicited action potentials, before drug. **E,** Confocal image (scale bar, 1000 μm) from one rat showing retro-Cre expression in the GPe. **F,** Confocal image (scale bar, 100μm) from one rat showing DIO-hM3D expression in the pDMS. **G,** Schematic showing the extent of retro-Cre spread in the GPe, imaged at the widest point for all included animals (top) and the extent of DIO-hM3D expression, imaged at the widest point for each rat (bottom). **H,** Total number of mCherry positive cells/mm2 in the pDMS in rats from each experimental group. **I,** Summary of experimental groups; blue arrows represent direct pathway and red arrows represent indirect pathway, thicker arrows indicate increased activity and thinner arrows indicate decreased activity. **J,** Mean (± SEM) lever presses per minute averaged across each day of instrumental training for each group. **K,** Mean lever presses per minute on the devalued and valued lever for each rat in each group, averaged across two days of non-rewarded choice tests.

Next, we assessed the functional consequence of iSPN stimulation on goal-directed learning using the same behavioural design described previously (Figure 2L). Expression of retro-Cre in the GPe and DIO-hM3D in the pDMS is shown in Figure 3E and 3F, respectively, and the location and spread of each injection for all rats is shown in Figure 3G. The mean number of cells expressing DIO-mCherry or DIO-hM3D was quantified for all animals, and is presented in Figure 3H; there were no significant differences between groups or viruses (Fs<2.6). There were three groups of rats, summarized in Figure 3I: all groups received retro-Cre into the GPe, and there were two control groups; one expressed DIO-mCherry on iSPNs and received CNO injections throughout training and the other expressed DIO-hM3D on iSPNs and received VEH injections throughout training. The third group expressed DIO-hM3D on iSPNs and received CNO injections throughout training to stimulate the activity of iSPNs in the pDMS. We trained rats in the same manner as described previously (Figure 2L). We found no effect of stimulating the activity of iSPNs during instrumental training on press rates (Figure 3J; F<1), and found that this stimulation did not affect goal-directed learning (Figure 3K; main effect of devaluation, (F(1,17)=34.561, *p*<0.001; no devaluation x group interaction, Fs<1.5). Therefore, neither inhibition nor stimulation of iSPNs during instrumental training produced any detectable effect on goal-directed learning.

### Bilateral optogenetic inhibition of dSPNs or iSPNs does not affect the performance of goal-directed actions

Having established that goal-directed learning is dependent on the activity of dSPNs but not iSPNs, we next sought to assess the involvement of these pathways in the performance of goal-directed actions. We used the same dual-virus approach described above to express halorhodopsin (AAV5-EF1a-DIO-eNpHR3.0-eYF; DIO-eNpHR) on dSPNs (Figure 4A) or iSPNs (Figure 4B), to selectively inhibit each pathway using an orange LED delivered to the pDMS. Retro-Cre was injected into the SNr (Figure 4C) or GPe (Figure 4D) to target dSPNs or iSPNs, respectively, and all rats were injected with DIO-eNpHR (Figure 4E-F) or a control fluorophore (AAV5-EF1a-DIO-eYFP; DIO-eYFP) into the pDMS (see Figure 4G). Groups used in this experiment are summarized in Figure 4H. Rats in the eYFP control group had DIO-eYFP in either dSPNs or iSPNs. Due to the lack of eNpHR construct, this group of rats retained intact pathway function and, indeed, as there were no differences in their press rates during training and no differential effects of LED light presentation on test (Fs<1), they were pooled into a single control group.

**Figure 4.**
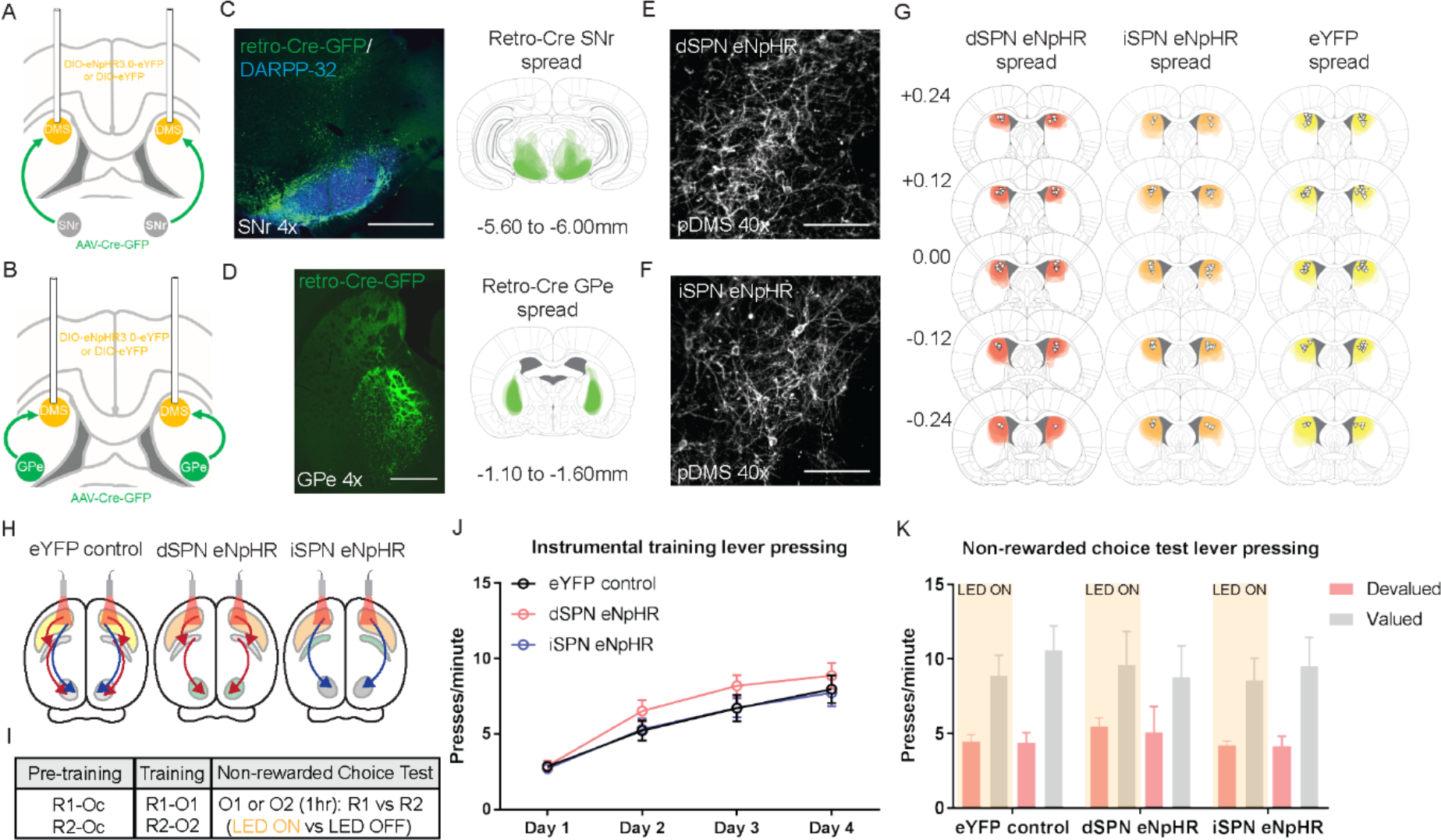
Bilateral optogenetic inhibition of dSPNs or iSPNs leaves the expression of goal-directed learning intact. **A,** Schematic depicting the surgery design used to target dSPNs; Retro-Cre was injected bilaterally into the SNr and DIO-eNpHR or DIO-eYFP was injected bilaterally into the pDMS. Cannulae were inserted bilaterally into the pDMS. **B,** Same as A but for targeting iSPNs; retro-Cre was injected into the GPe. **C,** Confocal image (scale bar, 1000 μm) from one rat showing retro-Cre expression in the SNr (left) and the extent of retro-Cre spread in the SNr, imaged at the widest point for all included animals (right). **D,** Same as C but showing retro-Cre expression in the GPe. **E,** Confocal image (scale bar, 100μm) from one rat showing DIO-eNpHR expression in pDMS dSPNs. **F,** Same as E but showing DIO-eNpHR expression in pDMS iSPNs. **G,** Extent of DIO-eNpHR and DIO-eYFP spread for rats in each group, imaged at 5 different AP locations encompassing where virus expression is at its widest. Triangles represent locations of cannula placements for each rat. **H,** Summary of experimental groups; blue arrows indicate intact direct pathway function and red arrows indicate intact indirect pathway function. **I,** Summary of the experimental design; R1 and R2 indicate left and right lever responses; Oc, O1 and O2 indicate distinct food outcomes; LED ON vs LED OFF indicates training or test days when LED light was delivered. **J,** Mean (± SEM) lever presses per minute averaged across each day of instrumental training for each group. **K,** Mean (± SEM) total number of lever presses on the devalued and valued lever in each group, averaged across two days of non-rewarded choice tests during LED ON (orange shaded) and LED OFF (non-shaded) periods.

A summary of the behavioural design is shown in Figure 4I. Instrumental pre-training and training sessions were conducted without LED light delivery. All groups acquired the instrumental response across pre-training (Figure S2A). Across four days of instrumental training (Figure 4J), all groups increased their press rates (F(2,43)=82.966, *p*<0.001) and there were no differences between groups in overall response rates (all Fs<1). We tested for goal-directed action using outcome devaluation followed by a choice extinction test. For the test, dSPNs or iSPNs in pDMS were inhibited with orange LED light during the first and last 2.5 minutes, separated by 2.5 minutes with no LED light. Responding in the LED ON periods was similar (all Fs<1) and so averaged for comparison with the LED OFF period. We found a main effect of devaluation (F(1,22)=37.568, *p*<0.001 – Figure 4K) but no effect of LED and no group x LED x lever interaction (Fs<1), indicating that rats showed intact outcome devaluation, and this was unaffected by optogenetic inhibition of either dSPNs or iSPNs. Additionally, optogenetic inhibition of dSPNs or iSPNs had no effect on the rats’ ability to discriminate between outcomes or levers during a rewarded devaluation test (Figure S2B; main effect of lever (valued vs devalued) F(1,22)=13.783, *p*=0.001; no group x lever, group x LED or group x LED x lever interactions, Fs<2.9). Therefore, unlike goal-directed learning, bilateral optogenetic inhibition of dSPNs had no effect on goal-directed performance.

### Unilateral inhibition of dSPNs, but not iSPNs, in the pDMS induces an ipsilateral response bias

To further examine the role of dSPNs and iSPNs in performance we used the same animals and examined the effect of unilateral suppression of these SPNs on response bias in a choice extinction test without devaluation (i.e. without pre-feeding; – Figure 5A). As represented in Figure 5B, a single patch cord was attached to either the left or right cannula (first and second test, respectively) and rats were again tested for 7.5 minutes as described previously. Again there were no differences between the first and last LED ON periods and so these blocks were pooled and compared to the LED OFF block. Test data are presented in Figure 5C. Although we again observed no main effects of press rate, lever side or LED, we now observed a significant group x lever x LED interaction (F(2,22)=3.456, *p*<0.05), indicating that the effect of the light on lever choice differed across groups. To further assess this, we conducted follow up tests in each group separately to assess whether rats’ response bias was affected by LED, and found that, whereas there were no significant interactions in Group iSPN eNpHR or Group eYFP control (Fs<1), the lever x LED interaction was significant for Group dSPN eNpHR (F(1,5)=8.924, *p*=0.031); i.e., the lever bias during the LED ON period favoured the ipsilateral lever, and this was reversed during the LED OFF period. We also analysed this data by comparing responding on the ipsilateral or contralateral lever during the LED ON period as a ratio of the total responding (LED ON plus LED OFF periods) for each lever (Figure 5D). For rats in Group iSPN eNpHR and Group eYFP, responses on each lever during the LED ON period were ~50% of the total responding on each lever, whereas for rats in Group dSPN eNpHR we observed a significant ipsilateral bias, (F(1,5)=10.342, *p*=0.024); 59% vs. 45% for ipsilateral vs. contralateral levers, respectively. Therefore, although, like bilateral inhibition, unilateral inhibition of dSPNs and iSPNs had no overall effect on performance, unilateral dSPN inhibition but not iSPN inhibition produced an ipsilateral bias in goal-directed performance.

**Figure 5.**
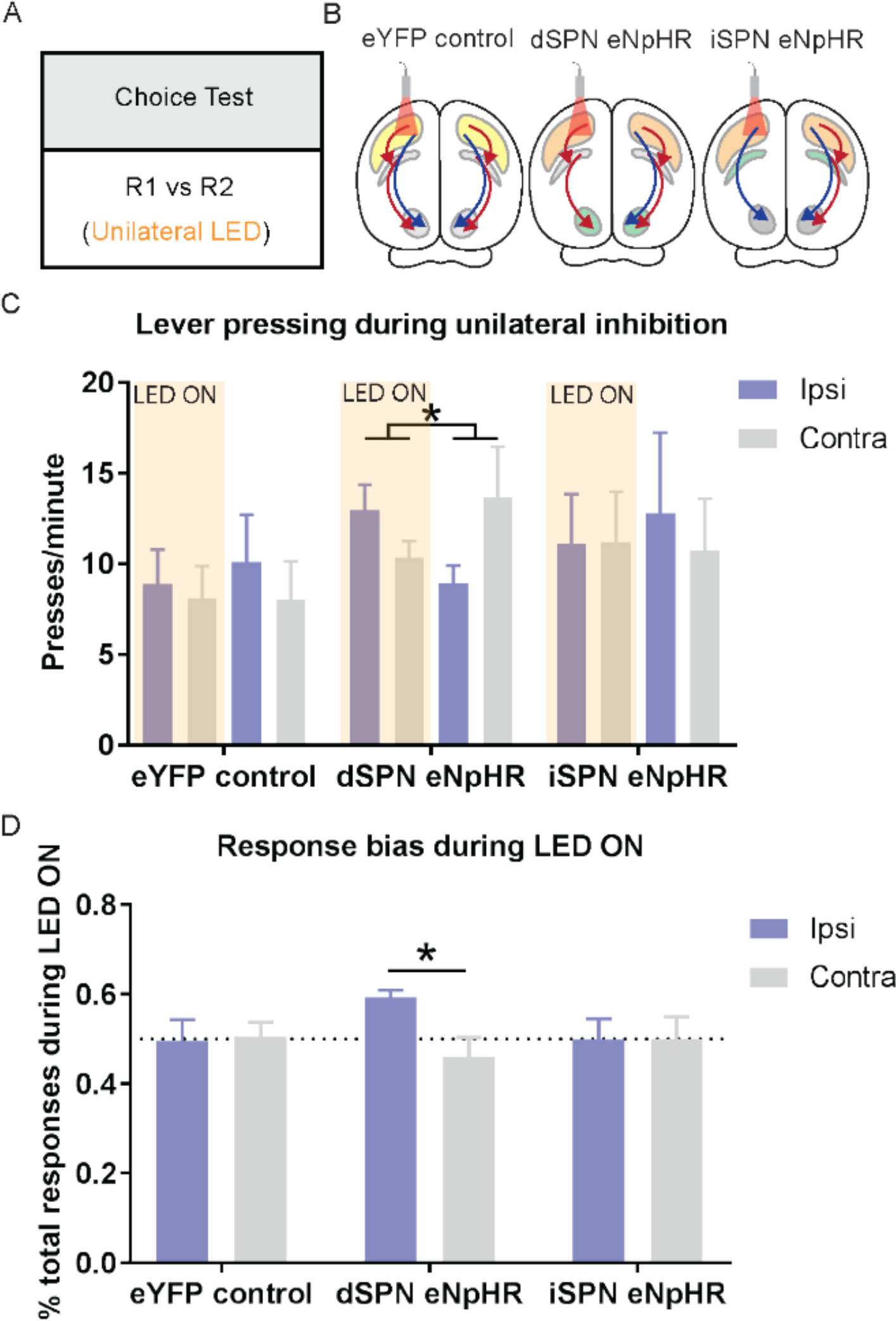
Unilateral inhibition of dSPNs, but not iSPNs, in the pDMS induces an ipsilateral response bias. **A,** Summary of the experimental design; R1 and R2 indicate left and right lever responses; rats were tested with unilateral LED light delivery. **B,** Summary of experimental groups; blue arrows indicate intact direct pathway function and red arrows indicate intact indirect pathway function. **C,** Mean (± SEM) presses per minute on the lever ipsilateral and contralateral to unilateral optogenetic inhibition in all groups during periods of LED ON (orange shaded) and LED OFF (non-shaded), averaged across two tests with inhibition in each hemisphere. **D,** Mean (± SEM) proportion of responding on the ipsilateral and contralateral lever during the LED ON period, as a proportion of the total responding on each respective lever, for all groups, averaged across two tests \**p* < 0.05.

### Optogenetic inhibition of iSPNs, but not dSPNs, in the pDMS impairs updating the action-outcome contingency

Our experiments so far have failed to detect any significant role for iSPNs in the acquisition and performance of goal-directed actions. In fact, recent findings suggest that iSPN activity may only be necessary for learning (and so performance) when instrumental contingencies change (Matamales *et al.*, 2020). In order to assess this possibility, we examined the effect of dSPN and iSPN inhibition when action-outcome contingencies were altered by a series of reversals in a two-choice situation. We used an instrumental reward reversal paradigm, wherein rats were exposed to the same action-outcome contingencies as during initial training, but with only one lever rewarded at a time and the other lever non-rewarded (i.e., if R1-O1 then R2-Ø; whereas if R2-O2 then R1-Ø). These contingencies were reversed every 2.5 min in semi-random fashion (no more than two of the same trial type sequentially). Each test contained 10 × 2.5 min trials, and rats were tested each day for 3 days. LED light was delivered bilaterally during all trials, but not between trials (Figure 6A-B).

**Figure 6.**
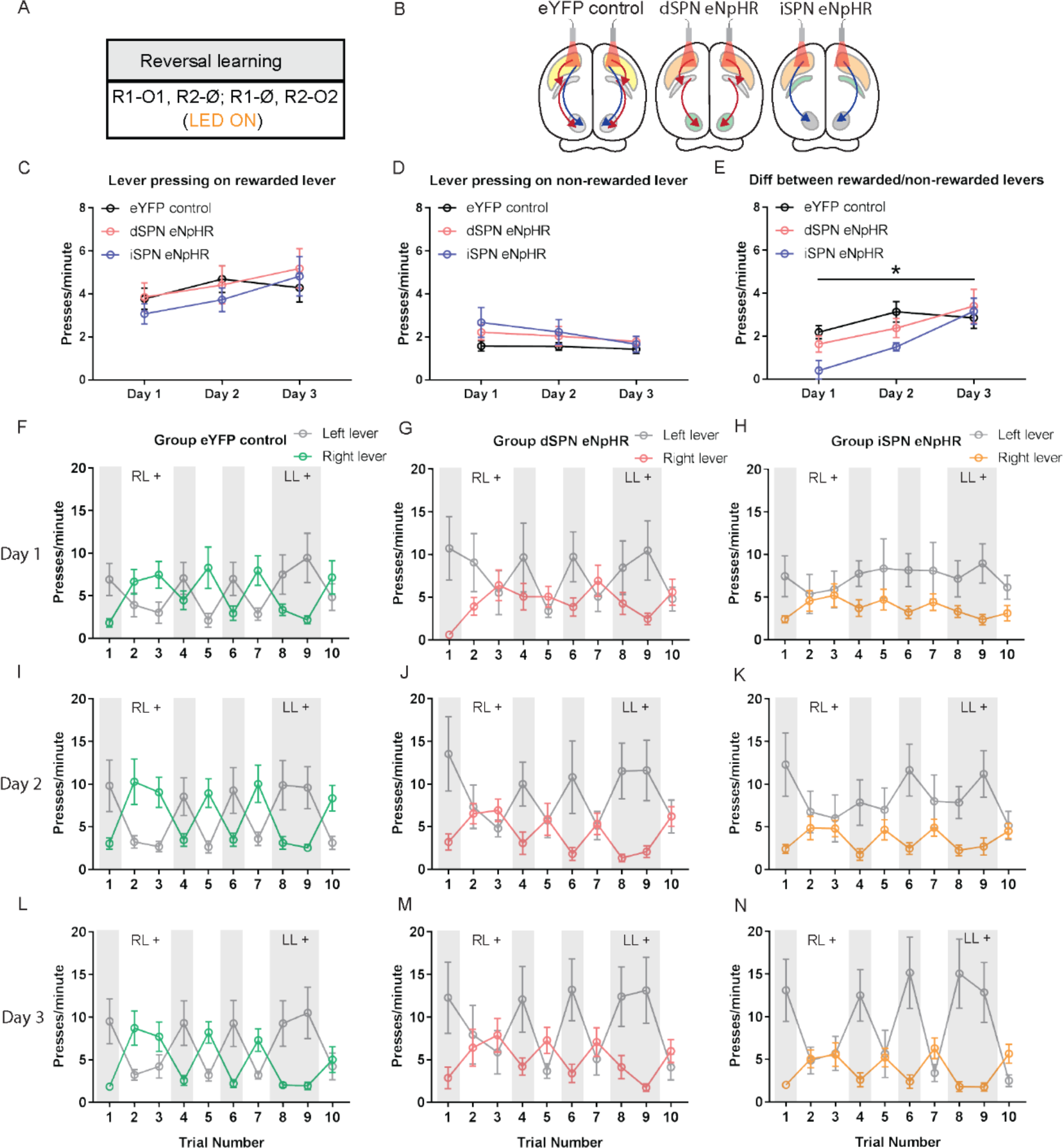
Optogenetic inhibition of iSPNs, but not dSPNs, in the pDMS impairs response flexibility. **A,** Summary of the experimental design; R1 and R2 indicate left and right lever responses; O1 and O2 indicate distinct food outcomes; Ø indicates non-rewarded responses; LED light was delivered during all lever presentations. **B,** Summary of experimental groups; blue arrows indicate intact direct pathway function and red arrows indicate intact indirect pathway function. **C,** Mean (± SEM) presses per minute on the rewarded lever across 3 days of reversal training in all groups. **D,** Mean (± SEM) presses per minute on the non-rewarded lever across 3 days of reversal training in all groups. **E,** Mean (± SEM) difference in press rate (expressed as presses per minute) between the rewarded and non-rewarded levers for each group. **F-H,** Mean (± SEM) lever presses per minute on the left lever and right lever for rats in each group on Day 1 of training, averaged across each 2.5-minute trial - grey shaded regions indicate trials in which the left lever was rewarded (LL+) and non-shaded regions indicate trials in which the right lever was rewarded (RL+). **I-K,** Same as previous but for Day 2. **L-N,** Same as previous but for Day 3. \**p*<0.05.

Press rates on the rewarded lever, the non-rewarded lever, and the difference in press rates between the two levers is presented in Figures 6C-E. All groups increased responding on the rewarded lever (Figure 6C; F(1,22)=11.6, *p*=0.003) and decreased responding on the non-rewarded lever across days (Figure 6D; F(1,22)=9.8, *p*=0.005). We analysed the difference in press rates between the rewarded and non-rewarded levers: The magnitude of this difference increased across training days (linear trend F(1,22)=19.8, *p*<0.001). There was no difference between Groups dSPN eNpHR and eYFP control (F<1), but there was a significant difference between Groups iSPN eNpHR and eYFP control (F(1,22)=4.4), and a significant Group (iSPN eNpHR vs eYFP control) x Day (linear) interaction, indicating that the difference between these groups was greater on Day 1 of training than on Day 3 (F(1, 22)=5.7, *p*=0.026).

Figure 6F-N shows the rate of responding on the left and right levers for each group, across three days of training – shaded blocks indicate those in which the left lever was rewarded and unshaded those in which the right lever was rewarded. We analysed each day separately: Rats in Group eYFP control and dSPN eNpHR updated performance according to the positive contingency during each epoch on Day 1 (Figure 6F-H); however, rats in Group iSPN eNpHR failed to update their performance as the positive contingency shifted across epochs. Supporting this, we found a significant lever x trial x group interaction between Groups iSPN eNpHR and eYFP controls (Bonferroni, k=3, F(1,22)=12.3, *p*=0.002) indicating that Group iSPN eNpHR were impaired at updating performance on Day 1. This pattern was maintained on Day 2 (Figure 6I-K): There was a significant main effect of reversal (lever x trial, Bonferroni, k=3, F(1,22)=87.3); this didn’t differ between Groups dSPN eNpHR and eYFP control (Bonferroni, k=3, F(1,22)=1.5), but there was a significant Group x reversal interaction between Groups iSPN eNpHR and eYFP control (Bonferroni, k=3, F(1,22)=8.4, *p*= 0.008). Finally, on Day 3 (Figure 6L-N), groups maintained their sensitivity to contingency (main effect of reversal, Bonferroni, k=3, F(1,22)=80.5, *p*<0.001), and there were no differences in the magnitude of reversal between Groups dSPN eNpHR and eYFP control (F<1), nor between Group iSPN eNpHR and eYFP control (F<1). Therefore, optogenetic inhibition of iSPNs impaired rats’ ability to flexibly adjust their instrumental performance in accordance with changes in the action-outcome contingency during the first two days of reversal learning consistent with the claim that iSPNs are necessary to update instrumental learning.

## Discussion

Using a novel, target-based approach to examine the involvement of the direct and indirect pathway projection neurons in goal-directed learning, performance, and action updating, we found evidence that goal-directed learning recruits dSPNs, but not iSPNs, that the role of this recruitment for performance is specific to the orientation of the response in contralateral space, and, as a consequence, that unilateral inhibition of dSPNs, but not iSPNs, produced an ipsilateral response bias. Furthermore, and most importantly, despite a lack of involvement in initial learning and performance, we found that iSPNs, but not dSPNs, were necessary for rapidly updating goal-directed learning when the action-outcome contingency changes to support response flexibility.

### SPNs and Goal-directed Learning: A Specific Function of the Direct Pathway

Prior research focused on assessing the involvement of dSPNs and iSPNs in goal-directed learning has generally inferred a greater relative involvement of dSPNs based on the use of cell-type specific GFP expression in either D1-GFP (Maroteaux *et al.*, 2014; Shan *et al.*, 2014) or D2-GFP (Shan *et al.*, 2014; Matamales *et al.*, 2020) mice. We confirmed this involvement using the target-based approach described here, both in the recruitment of Zif268 signalling, which was specifically increased in dSPNs and not iSPNs following instrumental training, and using chemogenetic inhibition, where we found causal evidence that dSPNs but not iSPNs are required for initial goal-directed learning. Furthermore, this effect of dSPN inhibition on goal-directed learning was specific to corticostriatal transmission at dSPNs, and was not due to changes in the relative balance between the two populations of pDMS projection neurons; chemogenetic stimulation of iSPNs during the same period of training had no effect on goal-directed learning.

In a functional sense, previous studies conducting on-line assessments of instrumental conditioning have told us several things regarding the role of SPNs in instrumental performance (e.g. Tecuapetla *et al.*, 2016) and even choice bias in relation to reward history (e.g. Tai *et al.*, 2012). However, what such assessments cannot determine is whether the actions being observed are in fact goal-directed. In order to make that assertion, tests are required to demonstrate that subjects have encoded the relationship between actions and their consequences, and are sensitive to the current value of the outcome (Balleine, 2019). Tests for outcome devaluation under extinction conditions fulfil both criteria; when animals bias responding away from a devalued lever, they demonstrate awareness of the action-outcome contingencies, and the capacity to modulate their choice based on the current value of each outcome in the absence of any feedback to inform this choice (Balleine & Dickinson, 1998). Importantly, the current experiments established the functional role of both dSPNs and iSPNs in goal-directed learning for actions verified to be goal-directed.

### SPNs and Goal-directed Performance: Involvement of the Direct Pathway

We have previously argued (Peak *et al.*, 2019) that glutmatergic inputs to the dorsal striatum constitute “learning” pathways in the broader basal ganglia circuitry, and striatal output pathways constitute “performance” pathways. Central to that argument is that the dorsal striatum is a point of convergence of these two functions and the current results refine this, suggesting that both goal-directed learning and performance relies specifically on dSPN input and ouptut processes in the pDMS.

While training-induced elevations of dSPN signalling have been reported, here we assessed each hemisphere separately, relative to the position of the lever and found that this is elevation was response-specific; i.e., it was lateralized with respect to the trained response. There is reason to predict this involvement is primarily driven by performance factors. For example, the activity of dopamine neurons projecting from the SNc to the DMS is lateralized according to response type; when rats press a left lever, SNc-DMS dopamine neurons in the right hemisphere show increased activity, and this is independent of action value (Parker *et al.*, 2016; Lee *et al.*, 2019). Therefore, we predict that lateralisation of activity in dSPNs during goal-directed performance is driven by lateralized dopamine input converging with a bilateral input from prefrontal cortex (Hart *et al.*, 2018a, 2018b).

Functional studies investigating the role of DMS SPNs in the performance of goal-directed actions are limited, and their findings have been contradictory. For example, Poyraz *et al.* (2016) found that DMS iSPN inhibition energizes action initiation, but does not affect performance of a goal-directed action tested under extinction. By contrast, Tai *et al.* (2012) reported that unilateral optogenetic stimulation of iSPNs biased instrumental responding towards the ipsilateral lever (or away from the contralateral lever). As we have argued, targeting these pathways using receptor expression as a proxy for each pathway (D2 or A2A receptors in iSPNs), is likely to have significant effects on how pathway function is interpreted. A second source of contradiction lies in the method of testing; when rats are tested under extinction (as in Poyraz *et al.*, 2016), they must rely on previously learned associations to guide performance, whereas, when outcomes continue to be delivered (as in Tai *et al.*, 2012), performance can be driven by either prior learning or new learning, conflating instrumental learning and performance effects. Here, we assessed the role of dSPNs and iSPNs on goal-directed performance under extinction, forcing animals to rely on prior learning.

We found that bilateral optogenetic inhibition, whether of dSPNs or iSPNs, at test did not affect goal-directed performance. Nevertheless, in the same animals, we found that unilateral optogenetic inhibition of dSPNs was effective in inducing an ipsilateral response bias (relative to the hemisphere of inhibition) in goal-directed responding, whereas unilateral iSPN inhibition had no effect. This effect of unilateral, but not bilateral inhibition of dSPNs on goal-directed performance emphasizes the importance of lateralized dSPN activity in inducing this response bias. By bilaterally dampening dSPN output, we did nothing to the relative balance of dSPN activity in each hemisphere, and as such, any existing response bias was unchanged. Lateralized control of motor output by dSPNs has been well demonstrated. SPN activity has been found to precede movement initiation and to predict the occurrence of contraversive movements in a lever pressing task (Cui *et al.*, 2013). Furthermore, unilateral activation or suppression biases contralateral or ipsilateral movements, respectively (Kravitz *et al.*, 2010; Bay Konig *et al.*, 2019), whereas unilateral stimulation of dSPNs produces a contralateral bias in lever pressing (Tai *et al.*, 2012). Our results indicate that these findings extend to goal-directed actions, consistent with the finding that DA terminals in the DMS respond more strongly during contralateral choices than ipsilateral choices and that this activity, too, precedes action performance (Parker *et al.*, 2016; Lee *et al.*, 2019).

### SPNs and Goal-directed Updating: A Role for the Indirect Pathway

Despite a lack of involvement in initial learning, we found that bilateral optogenetic inhibition of pDMS iSPNs, but not dSPNs, impaired the updating of goal-directed learning; specifically, the contingency controlling the goal-directed actions of rats exposed to a series of reversals. Rats that had iSPNs inhibited were slower to learn which of two levers was rewarded and to switch their responding appropriately. Importantly, this failure was overcome with continued training, suggesting that with enough training, rats are capable of learning response strategies to compensate for this impairment. A lack of flexibility in this task could be due to several factors, but primarily, impairments can manifest as an inability to suppress responding on the non-rewarded lever (perseveration), or a failure to increase responding on the rewarded lever. Although we found no significant effect on either of these factors alone, the overall impairment by rats with iSPN inhibition appears to be due to their combination. Indeed, there is recent evidence to suggest that both of these forms of response flexibility require iSPNs; Matamales *et al.* (2020) demonstrated that ablating iSPNs in the DMS encourages recurrent responding during the extinction of goal-directed learning (response perseveration), and Nonomura *et al.* (2018) have shown that iSPNs in the DMS encode non-rewarded responses, and, if optogenetically stimulated following a non-rewarded response, promote switching to an alternate response.

Although it might be supposed that dips in phasic dopamine activity that occur when outcomes are omitted could form the basis for iSPN involvement in these forms of reversal, or in extinction learning, Matamales *et al.* (2020) found similar involvement of iSPNs when animals were exposed to reversal in the identity of the outcomes of two other continuously rewarded actions. It seems reasonable to conclude, therefore, that the causal evidence for the involvement of iSPNs in contingency reversal observed in the current study supports the involvement of iSPNs in action-outcome updating rather than action suppression, guiding changes in plasticity associated with non-reward, and interleaving that learning with prior encoding so as to minimize interference between them.

## Conclusions

The control of instrumental actions according to our current goals requires the recruitment of dSPN and iSPN pathways differentially; by assessing each pathway according to projection target, rather than receptor expression, we were able to dissociate their functions in goal-directed actions along several lines. Broadly, dSPNs are important for encoding goal-directed learning whereas goal-directed performance is reflected in, and modulated by, asymmetrical activity in dSPNs in each hemisphere driving contralateral responses. By contrast, when contingencies change, iSPNs are required to update those contingencies to support response flexibility.

## METHODS

### SUBJECT DETAILS

Subjects for all experiments were Long-Evans rats, obtained either from the Animal Resource Centre (Perth, Western Australia) or from the Randwick Rat Breeding Facility (Randwick, New South Wales). All rats were healthy and experimentally naïve at the beginning of each experiment and at least 8 weeks old prior to surgery. Rats were housed in transparent, plastic boxes with 2-4 rats per box in a climate-controlled colony room and maintained on a 12 hr light/dark cycle (lights on at 0700). All experiments were conducted during the light phase. Water and standard lab chow were available ad libitum prior to the start of experiments. For all experiments, rats were randomly assigned to experimental groups. All experiments conformed to the guidelines on the ethical use of animals maintained by the *Australian code for the care and use of animals for scientific purposes*, and all procedures were approved by the Animal Care and Ethics Committee at either the University of New South Wales or the University of Sydney. Specific subject details for each experiment are provided below.

#### SPN tracing and immunofluorescence experiment

Subjects were 25 experimentally naïve female outbred Long-Evans rats bred within the laboratory from animals obtained from the Animal Resource Centre (Perth, Western Australia).

#### Electrophysiology recordings

Subjects for recordings of SPNs were 7 (4 for dSPNs and 3 for iSPNs) experimentally naïve female Long-Evans rats obtained from the Randwick Rat Breeding Facility (Randwick, New South Wales).

#### DREADDs suppression experiment

Subjects were 79 experimentally naïve Long-Evans rats (36 male and 43 female) obtained from either the Animal Resource Centre (Perth, Western Australia) or from the Randwick Rat Breeding Facility (Randwick, New South Wales).

#### DREADDs stimulation experiment

Subjects were 30 experimentally naïve Long-Evans rats (14 male and 16 female) obtained from the Randwick Rat Breeding Facility (Randwick, New South Wales).

#### Optogenetic inhibition experiment

Subjects were 39 experimentally naïve Long-Evans rats (13 male and 24 female) obtained from the Randwick Rat Breeding Facility (Randwick, New South Wales).

### METHOD DETAILS

#### Surgery

Rats received stereotaxic surgery conducted under isoflurane gas anaesthesia (0.6L/min; isoflurane at 5% during induction and 2-2.5% maintenance). Animals were placed in a stereotaxic frame (Kopf instruments) and a subcutaneous injection of bupivacaine hydrochloride at the incision site and a subcutaneous injection of carprofen in the lower flank administered. An incision was made to expose the scalp, membrane on top of skull was cleared and the head was adjusted to align bregma and lambda on the same horizontal plane. Small holes were drilled in the skull above the target structures and a Hamilton syringe (1.0μl) connected to an Infuse/Withdraw pump (Harvard Apparatus) or a capillary micropipette connected to a Nanonject (Drummond Scientific) was lowered into the brain for infusions of viruses or tracers, respectively. Following each infusion, the needle was left in position for 3-5 minutes to allow diffusion. Following the final infusion, the incision site was closed and secured with surgical staples. Rats then received an injection of antibiotic (either Benacillin (0.3ml i.p.) or Duplocillin (0.15ml/kg s.c.)) and saline (3ml i.p.). Rats were given a minimum of 2 weeks recovery time following surgery to allow for sufficient tracer expression, or 4 weeks for viral expression.

Surgical co-ordinates for each region were pre-determined from pilot studies and varied slightly between experiments. All infusions were bilateral. Retrograde tracers or retro-Cre were infused into the GPe or SNr at the following coordinates (mm from bregma): males: GPe: AP −1.25; ML ±3.1; DV −6.9; SNr: AP −5.8; ML ±1.95; DV −8.8, females: GPe: AP −1.1; ML ±3.0; DV −6.9; SNr: AP −5.6 to −5.8; ML ±1.9; DV −8.6 to −8.65. Channelrhodopsin was infused into the PL at the coordinates: AP +2.9; ML ±0.6; DV −3.8. DREADDs, halorhodopsin or control fluorophores were infused into the pDMS at the following coordinates: males: AP −0.3; ML ±2.35 to ±2.40; DV −4.7, females, AP −0.25; ML ±2.3 to ±2.35; DV −4.5 to −4.6. For rats in optogenetic inhibition experiment, glass cannulae were inserted above the pDMS at the following coordinates: males: AP −0.3; ML ±2.40 to ±2.45; DV −4.50, females: AP −0.25; ML ±2.35 to ±2.40; DV −4.35 to −4.40.

#### Viruses and Tracers

SPNs were targeted by infusing the retrograde tracers Cholera-Toxin B (CTB; 1% in dH2O; List Biological Laboratories) and Fluorogold (FG; 3% in saline; Fluorochrome) into the GPe and SNr, respectively. Both tracers were infused at a total volume of 82.2nl and a rate of 23nl/minute. Volumes were selected based on pilot studies designed to contain the spread of tracers within each region.

Projection neurons in the PL were targeted by infusing Channelrhodopsin, AAV5-CaMKIIa-hChR2(H134R)-EYFP (ChR2-eYFP) into the PL at a total volume of 300nl and a rate of 150nl/minute.

Striatal SPNs were targeted by infusing retro-Cre (rAAV5.CMV.HI.eGFP-CRE.WPRE.SV40) into either the SNr (for dSPNs) or GPe (for iSPNs). Retro-Cre was infused into the SNr at a total volume of between 300nl and 500nl at a rate of 100nl/minute, and into the GPe at a total volume of 150nl and a rate of 50nl/minute.

All DREADDs viruses: DIO-hM4D (rAAV5/hSyn-DIO-hM4D(Gi)-mCherry), DIO-mCherry (rAAV5/hSyn-DIO-mCherry), and DIO-hM3D (rAAV5/hSyn-DIO-hM3D-mCherry), were infused into the pDMS at a total volume of 750nl at a rate of 100-150nl/minute.

For optogenetic inhibition, 500nl of DIO-eNpHR (AAV5-EF1a-DIO-eNpHR3.0-eYFP) or DIO-eYFP (AAV5-EF1a-DIO-eYFP) was infused into the pDMS at a rate of 100nl/minute.

For all virus and tracer infusions, the needle was left in place for a minimum of 3 minutes following each infusion.

#### Drugs

For DREADDs experiments, Clozapine-N-Oxide (CNO) was used as the ligand to activate either the hM4Di or hM3Dq DREADDs receptors. CNO was made on the morning of each required test day; dissolved in 0.6% 5M HCl and diluted with distilled water (dH2O). Vehicle consisted of 0.1% 5M HCl diluted with dH2O and pH was matched with that of the CNO in solution. For DIO-hM4D, CNO was made to a concentration of 7mg/mL and injected at a dose of 7mg/kg. For DIO-hM3D, CNO was made to a concentration of 3mg/mL and injected at a dose of 3mg/kg.

#### Apparatus

For immunofluorescence and DREADDs experiments, training and testing were conducted in 16 Med Associates operant chambers, individually housed in light and sound attenuating cabinets. Each chamber was fitted with a food magazine connected to a 45mg grain pellet (Bioserve Biotechnologies) dispenser as well as two pumps fitted with external syringes that delivered either 0.2ml of 20% sucrose solution (white sugar, Coles, Australia) or 0.2ml of 20% maltodextrin solution (Poly-Joule, Nutrica, Australia). An infrared detector was situated horizontally across the inside of the magazine to detect head entries.

The chambers were also fitted with two retractable levers, located on either side of the magazine. A house light (3W, 24V) was located in a central position at the top of the wall opposite to the food magazine and illuminated during all experimental stages, unless otherwise stated. Training and testing sessions were pre-programmed and controlled by computers external to testing rooms, using Med Associates software (Med-PC IV or V), which also recorded experimental data from each session.

For optogenetic experiments, LED light delivery was controlled through a TTL adapter, converting 28V DC output to a TTL transition (Med Associates, Georgia, Vermont). This was connected to an LED driver (Doric lenses) and Doric Connectorized LED light source with a fibre-optic rotary joint for connection to fibre optic patch cords. For all optogenetic experiments LED light stimulation was delivered as orange light (625nm) with a power of ~10mW measured at the cannula tip, pulsed at 40Hz.

For tests of goal-directed behaviour via sensory specific satiety, a separate devaluation room was used that contained sixteen individual, open top plastic boxes with stainless steel wire mesh lids. During devaluation, lights were kept off and each individual chamber was fitted with either a glass petri dish for pellet devaluation, or a plastic drink bottle with sipper for sucrose devaluation.

#### Behavioural Protocol and Food Restriction

##### Food restriction

For behavioural experiments, rats underwent 3 days of food restriction before the onset of magazine training. For the first two days of food restriction, male and female rats received 8 or 6 grams of standard lab chow, respectively, and this increased to 12 or 8 grams for the third day and remainder of experiment, adjusted dependent on rats’ weight across days. Their weight was monitored closely to ensure that their food restricted body weight did not fall below 85% of their free-feeding body weight.

##### Magazine training

Rats were given 2 days of magazine training during which the to-be-trained food rewards (either 45mg grain pellets and 0.2ml of 20% sucrose solution or 0.2ml of 20% polycose solution) was delivered at a random interval schedule of 60 seconds.

##### Instrumental pre-training

For experiments that included pre-training, rats received two daily sessions (one session on each of left and right lever, order counterbalanced within groups and across days) of lever press training each day. Lever press responses were rewarded with 0.2ml of 20% polycose solution, which on days 1 and 2 were delivered on a continuous reinforcement schedule (CRF; every lever press earned one reward). On day 3, lever presses were rewarded on a random interval 15 (RI15; presses were rewarded on average every 15 seconds). On days 5 and 6, lever presses were rewarded on a random interval 30 (RI30; presses were rewarded on average every 30 seconds). Criteria for progression from CRF on to RI15 training was 30 outcomes earned in a single session for each lever.

##### Instrumental training

*For SPN tracing with immunofluorescence experiment*; rats did not receive pre-training and instead received instrumental training sessions following two days of magazine training. Rats were trained on either the left or right lever, counterbalanced within groups. Lever press responses were rewarded at increasing intervals over training days. Training sessions progressed from CRF (days 1 and 2) to RI15 (day 3) to RI30 (days 5 and 6). *For DREADDs experiments;* following instrumental pre-training rats received two days of instrumental training, during which they received alternating presentations of each lever, which delivered separate outcomes (sucrose solution or grain pellets). Rats were presented with one lever (left or right, counterbalanced within groups), which earned one outcome (either pellets or sucrose, counterbalanced within groups) on an RI30 schedule of reinforcement. This lever was presented for either 10 minutes (dSPN rats; average 15.1 outcomes) or 15 outcomes (iSPN rats; average 9.2 minutes), after which it retracted, and the house light went off for a 1 min intertrial interval (ITI), and then the alternate lever was presented with the same criteria. This sequence was repeated, such that rats all received two presentations of each lever. *For optogenetic inhibition experiment*; instrumental training was as described for DREADDs experiments, however rats received four days of instrumental training on an RI30 schedule, with levers presented for 15 outcomes.

##### Outcome Devaluation

All rats were habituated to devaluation chambers with two 1-hour pre-exposure sessions conducted during the pre-training phase. On test days, rats were placed in devaluation chambers for 1 hour where they had ad libitum access to one of the previously earned outcomes, either sucrose solution or grain pellets. Rats were then immediately returned to the operant chambers for the choice test. Devaluation was conducted across two consecutive days of choice testing (for rewarded and non-rewarded choice tests), with one outcome (pellets or sucrose solution) devalued each day, order counterbalanced within groups.

##### Non-rewarded Choice Test

For *DREADDs experiments*, non-rewarded choice tests were conducted drug-free. Sessions began with the illumination of the house light, and simultaneous presentation of both left and right levers. No outcomes were delivered during this session and responses on left and right levers recorded, as well as magazine entries. Sessions ended after 5 minutes. For *optogenetic inhibition experiment*, non-rewarded choice tests were the same as described for DREADDs experiments, but lasted 7.5 minutes, during which orange LED light was delivered bilaterally into the brain in an ABA design with LED illumination on for the first and final 2.5 minutes of testing, separated by 2.5 minutes of no LED illumination.

##### Rewarded Choice Test

*For DREADDs experiments*, on rewarded choice test days, rats received an injection of CNO or vehicle prior to outcome devaluation and thus approximately one hour before test commencement. Both levers were rewarded with their previously earned outcomes (pellets and sucrose), on a random ratio 7 (RR7; on average every 7th press is rewarded) schedule, with the constraint that the first response on each lever was always rewarded. Sessions ended after 10 minutes. For *optogenetic inhibition experiment*, rewarded choice testing was as described for DREADDs experiments, however, each test lasted 7.5 minutes, with orange LED light delivered into the brain bilaterally for the first and last 2.5 minutes.

##### Non-rewarded Choice test with unilateral LED light delivery

Non-rewarded choice tests were conducted in the absence of devaluation pre-feeding and orange LED light delivered unilaterally into either the left (first test) and right (second test) hemisphere. This was done so that each rat received suppression to both hemispheres separately, and also so that for test 1 and 2, the outcome that was paired with the left or right lever, relative to instrumental training protocol, was counterbalanced within groups. Both tests occurred on the same day, approximately 1 hour apart for each rat. The test itself was identical to that described above for the non-rewarded choice test with bilateral LED light delivery.

##### Reversal training

Sessions began with the illumination of the house light, and after 1 minute, both the left and right levers were presented. Orange LED light was delivered into the brain for all periods when levers were presented. During lever presentation, only one lever was rewarded on an RI15 schedule of reinforcement (preserving the originally trained response-outcome pairings) and this alternated in an ABBABABAAB design whereby A corresponded to left lever rewarded and B corresponded to right lever rewarded. Each “block”, i.e. A or B, lasted for 2.5 minutes and was separated by a 1 minute inter-trial interval, during which both levers were retracted, but the house light remained illuminated. In this test, responding on the left and right levers were recorded during periods of lever presentation, as well as total magazine entries. Sessions ended after the 10th block (35 minutes) and rats were returned to their home cages.

##### Rotarod

An assessment of general motor coordination was conducted for all DREADDs experiments using the rotarod test. Rats received one day of rotarod training, drug-free, during which they received 3 trials on a fixed revolutions per minute (rpm) setting of 5rpm. Rats then received two days of rotarod testing on an accelerating mode, whereby the speed or rotation was set at a minimum of 5rpm, and this increased to 40rpm over a 4-minute period. On each of these test days rats received an injection of either CNO or vehicle, counterbalanced across rats. On each test day, rats received three trials and the best two trials were averaged to give a value for total time spent on the rotarod in seconds.

#### Histology

##### Transcardial perfusion

For tissue analysis, rats received an intraperitoneal injection of a lethal dose of pentobarbital (0.5-0.9ml) and were perfused transcardially with 400ml of cold 4% paraformaldehyde (PFA) in 0.1M phosphate buffer (PB). Fixed brains were immediately removed and stored in PFA for a further 12-72 hours. Brains were sliced into 40μm coronal sections in 0.1M phosphate buffer salt solution (PBS) on a vibratome (LeicaVT1000S) and stored at −30 degrees Celsius in a cryoprotective solution.

##### Immunofluorescence Protocol

Free-floating sections containing the region of interest were washed 3 times for 10 minutes each in 0.1M PBS solution. Sections were then incubated in a 0.5 Triton-X, 10% NHS in 0.1M PBS blocking solution for 2 hours. Following this, sections were transferred into a solution containing primary antibodies in 0.2% Triton-X, 2% NHS in 0.1M PBS for between 12 and 48 hours (dependent upon antibody) at either room temperature or 4 degrees Celsius. Sections were washed three times in 0.1M PBS for 10 minutes each and then incubated in a solution containing the secondary antibodies diluted in 0.2% Triton-X, 2% NHS in 0.1M PBS for 2 hours at room temperature. Sections were again washed three times in 0.1M PBS for ten minutes, and once in 0.1M PB for ten minutes and mounted onto glass slides using Fluoromount or Vectashield mounting medium. Sections that did not require immunofluorescence staining were washed 3 times for 10 min each in PBS and then for 10 min in PB and mounted in the same fashion. Images were generally taken the day following mounting using a confocal microscope (Olympus BX61WI) or a Spinning Disk Microscope (Andor Diskovery).

##### SPN tracing and immunofluorescence experiment

For simultaneous detection of DARPP-32, FG and CTB for tracing analysis, sections containing the pDMS were incubated with mouse anti-DARPP-32 (1:1000) and goat anti-CTB (1:2000). For simultaneous detection of Zif268, FG and CTB for activity analysis, sections containing the pDMS were incubated with rabbit anti-EGR1/Zif268 (1:300) and goat anti-CTB (1:2000). For placement analysis, sections containing the SNr and GPe were selected and immunostained for CTB (1:2000). Native fluorescence was used for identification of fluorogold throughout.

##### DREADDs experiments

Sections containing the pDMS were selected and virus expression imaged at the widest point. mCherry expression from DIO-hM4D, DIO-hM3D and DIO-mCherry were imaged without the addition of fluorescent antibodies. For placement analysis of retro-Cre, SNr sections were incubated with mouse anti-DARPP-32 (1:1000) and rabbit anti-GFP (1:1000) to label SNr boundaries and retro-Cre expression. GPe sections were incubated with rabbit anti-GFP (1:1000).

##### Optogenetic experiment

Five sections were selected that contained the pDMS at different anterior-posterior coordinates, and three selected that contained either the SNr (for dSPN targeting) or GPe (for iSPN targeting). pDMS sections were imaged for eYFP without the use of fluorescent antibodies. Verification of SNr and GPe placements of retro-Cre were the same as described for DREADDs experiments.

All immunofluorescence staining used secondary antibodies Alexa 488, Alexa 546 or Alexa 647 (1:1000).

#### Electrophysiology

##### Brain slice preparation

Rats were euthanized under deep anaesthesia (isoflurane 4% in air). Brains were rapidly removed and cut using a vibratome (Leica Microsystems VT1200S, Germany) in ice-cold oxygenated sucrose buffer containing (in mM): 241 sucrose, 28 NaHCO3, 11 glucose, 1.4 NaH2PO4, 3.3 KCl, 0.2 CaCl2, 7 MgCl2. Coronal brain slices (300 μm thick) containing the pDMS were sampled and maintained at 33ºC in a submerged chamber containing physiological saline with composition (in mM): 124 NaCl, 3.5 KCl, 1.25 NaH2PO4, 1 MgCl2, 1 CaCl2, 10 glucose and 26 NaHCO3, and equilibrated with 95% O2 and 5% CO2.

##### Electrophysiology recording

After equilibration for 1 h, slices were transferred to a recording chamber and neurons visualized under an upright microscope (BX50WI, Olympus, Shinjuku, Japan) using differential interference contrast (DIC) Dodt tube optics and mCherry fluorescence, and superfused continuously (1.5 ml/min) with oxygenated physiological saline at 33ºC. Whole-cell patch-clamp recordings were made using electrodes (2–5 MΩ) containing internal solution (in mM): 115 K gluconate, 20 NaCl, 1 MgCl2, 10 HEPES, 11 EGTA, 5 Mg-ATP, and 0.33 Na-GTP, pH 7.3, osmolarity 285-290 mOsm/L. Biocytin (0.1%) was added to the internal solution for marking the sampled neurons during recording. Data acquisition was performed with a Multiclamp 700B amplifier (Molecular Devices, Sunnyvale, CA), connected to a Macintosh computer and interface ITC-18 (Instrutech, Long Island, NY). Liquid junction potentials of −10 mV were not corrected. In current-clamp mode, membrane potentials were sampled at 5 kHz (low pass filter 2 kHz, Axograph X, Axograph, Berkeley, CA). Stock solution of drug was diluted to working concentration in the extracellular solution immediately before use and applied by continuous superfusion. Data from whole-cell recordings were only included in analyses if (1) the neurons appeared healthy under DIC on monitor screen, and (2) action potential amplitudes were at least 65 mV measured under current-clamp mode, to ensure that only highly viable neurons were included.

##### LED stimulation

For light-evoked terminal glutamate release impinged onto SPNs, ChR2-containing corticostriatal axons in the pDMS were stimulated by pulsing an LED light (473 nm, ThorLabs, New Jersey, LED light source attached to microscope) onto the slice under 40x water-immersion objective. Continuous trains (at 0.5 Hz) of light pulses (1ms width, interpulse interval 50 ms and 20 pulses per train) were delivered onto the slice at a maximum intensity of 3.9 mW to evoke action potentials arising from excitatory postsynaptic potentials. The stimulation protocol was chosen to mimic in vivo recordings in SPNs.

##### Post hoc histological analysis

Immediately after physiological recording, brain slices containing biocytin-filled neurons were fixed overnight in 4% paraformaldehyde/0.16 M phosphate buffer (PB) solution, rinsed and then placed in 0.3% Triton X-100/PB for 3 days to permeabilize cells. Slices were then incubated in AMCA-conjugated avidin (1:500; Vector Laboratory), or Streptavidin-conjugated Alexa 488 (1:500; Invitrogen), plus 2% horse serum and 0.2% Triton X-100/PB for 2 h to reveal biocytin-labelled neurons. Stained slices were rinsed, mounted onto glass slides, dried, and coverslipped with Fluoromount-G mounting medium (Southern Biotech). A 2D projection Image was later obtained from a collated image stack using confocal laser scanning microscopy (Fluoview FV1000, BX61WI microscope, Olympus).

### ANALYSES

#### Exclusions and group allocation

##### SPN tracing and immunofluorescence

Two rats were excluded from tracing analysis for misplaced injection sites leaving 23 rats. One other rat was excluded from activity marker quantifications due to damaged sections that were not quantifiable. This left 22 rats for analysis (Group Instrumental, n=11; Group Yoked, n=11).

##### DREADDs suppression experiment

Twenty-eight rats were excluded; 17 for low virus expression (<25 cells/mm2), 6 for virus spread beyond the DMS, 1 for infection, 3 that did not acquire the instrumental response and 1 that did not consume the outcome during devaluation pre-feeding. This left 51 rats for analysis, of which 23 were male and 28 were female (Group CNO control, n=13; Group dSPN+VEH, n=16; Group iSPN+VEH, n=4; dSPN+CNO, n=12; Group iSPN+CNO, n=6).

##### DREADDs stimulation experiment

Ten rats were excluded; 5 for low virus expression in the pDMS (<25 cells/mm^2^), and 5 for excessive virus spread not confined to the pDMS, leaving 20 rats for analysis, of which 11 were male and 9 were female (Group mCherry+CNO, n=6; Group hM4D+VEH, n=6; Group hM4D+CNO, n=8).

##### Optogenetic inhibition experiment

Fourteen rats were excluded; 2 for low virus expression in the pDMS, 11 for misplaced cannulae and 1 for infection, leaving 25 rats, of which 11 were male and 14 were female (Group eYFP control, n=11; Group dSPN eNpHR, n=6; Group iSPN eNpHR, n=8).

#### Immunofluorescence analysis

All immunofluorescence analysis was conducted using Image J software. Animals with misplaced injection sites or misplaced cannulae were excluded. Statistical analyses were performed using PSY software. The per-comparison error rate was controlled at alpha=0.05.

##### Activity marker

To maintain consistency for Zif268 quantification, all sections within a series were imaged within 3 days of immunofluorescence staining, and all images were taken within a 24 hour period. A brightness threshold was set for each series of sections that were immunostained and imaged together, and this was kept constant within a series. Only cells containing two or more pixels brighter than these values were counted. Cell counts were analyzed according to the total number of cells, calculated as the mean number of cells per mm^2^, per hemisphere, averaged across four slices per rat. Mean Zif268 was also calculated as a percentage of total dSPN or iSPN pathway; cells that were positive for Zif268 and either FG or CTB were expressed as a percentage of the total number of FG or CTB positive cells. Co-labelled cells, i.e. Zif268 + FG or Zif268 + CTB do not include counts for triple labelled cells, i.e. Zif268 + FG + CTB. Orthogonal contrasts were used to test for a main effect of group and orthogonal pairwise comparisons between each hemisphere in each group.

##### Retrograde tracing

For tracing analysis, cells that expressed FG, CTB and DARPP-32, as well as all co-labelled cells, were quantified and averaged across hemispheres, to give a mean number per mm^2^ for each hemisphere for each rat. Cells that were positive for both DARPP-32 and FG or CTB were represented as a percentage of total DARPP-32 positive neurons. Co-labelled cells, i.e. DARPP-32 + CTB or DARPP-32 + FG do not include counts for triple labelled cells, i.e. DARPP-32 + FG + CTB. Pairwise comparisons were used to test for differences in how many neurons were labelled with each retrograde tracer, FG or CTB and orthogonal contrasts were used to test for differences in the number of neurons that were labelled with each tracer in the pDMS in instrumental versus yoked rats.

##### DREADDs expression

Counts of mCherry positive cells in the pDMS were conducted manually with a cell counter in Image J and total number of cells expressed as cells/mm^2^. Orthogonal contrasts were used to compare the number of cells expressing DIO-hM4D between groups that expressed this virus on the same population of SPNs, i.e. dSPNs or iSPNs and to compare the number of cells infected with each virus (DIO-hM4D or DIO-mCherry).

Data from electrophysiological recordings were analayzed in GraphPad Prism 7. A paired t-test was used to compare the action potential frequency at baseline, to the action potential frequency after the application of CNO.

#### Behavioural analysis

For behavioural experiments, data collection was performed using Med-PC software (Versions IV and V), which was tabulated using Microsoft Excel and graphs generated with GraphPad Prism 7. Statistical analyses were performed using PSY software or IBM SPSS Statistics 26. The per-comparison error rate was controlled at α=0.05. For non-orthogonal contrasts, a Bonferroni correction was used to control the family-wise error rate at α=0.05

##### SPN activity marker experiment

Training data was analysed using orthogonal contrasts, testing for a main effect of group (instrumental versus yoked) and a linear trend analysis used to examine changes in magazine entries and press rates across training days.

##### DREADDs inhibition experiment

Non-orthogonal contrasts were used for all data analyses, and the family-wise error rate (FWER) controlled at α=0.05 using Bonferroni correction. All analyses tested for a main effect of group (dSPN+CNO versus controls, iSPN+CNO versus controls, Vehicle controls versus CNO control). For instrumental training, data was analyzed for a main effect of Group and training day, and any interactions. Non-rewarded choice test data was analysed for a main effect of Group and lever (devalued versus valued) and any interactions (Group x Lever). Rewarded choice test data was analysed for a main effect of Group and lever (devalued versus valued) and any interactions (Group x Lever). Rotarod data was analysed for a pairwise comparisons within each group, when rats were tested under CNO or vehicle.

##### DREADDs stimulation experiment

Planned orthogonal contrasts were used for all data analyses. Training data was analysed testing for a main effect of group (hM3D+CNO, versus controls, hM3D+VEH versus CNO control) and a main effect of training day. Non-rewarded choice test data was analysed testing for a main effect of group (as above) and a main effect of lever (devalued versus valued).

##### Optogenetic inhibition experiment

Training data was analysed using a two way repeated measures ANOVA F test for a main effect of Group and a main effect of training day. Choice test data (non-rewarded, rewarded and choice test with unilateral inhibition) were analysed using a three-way ANOVA F test for main effect of Group, period (LED ON or LED OFF) and lever (devalued or valued). Significant interactions were followed up by testing for the effect of LED light on choice in each group separately (significance determined by Scheffe). Reversal training data was analysed using planned contrasts testing for effects of Group (dSPN eNpHR or iSPN eNpHR vs eYFP control) and training day on pressing the rewarded or non-rewarded lever. Trial-by-trial reversal data were analysed separately for each group with non-orthogonal contrasts testing for a main effect of reversal (lever x trial type interaction) and reversal x group interactions across each day, controlling the FWER at α=0.05 using Bonferroni correction.

## Author contributions

JP, GH and BB designed the experiments, JP and BC performed the experiments, JP, GH, BC and BB wrote the manuscript.

## Acknowledgements

The research reported in this manuscript was supported by a Discovery Grant from the Australian Research Council, #DP150104878, and a both a Project Grant, #GNT 1165346, and a Senior Principal Research Fellowship, #GNT1079561, from the National Health and Medical Research Council of Australia to BWB. The authors thank J Bertran-Gonzalez for comments on the manuscript.

## Correspondence

Bernard Balleine, Decision Neuroscience Laboratory, School of Psychology, UNSW Sydney, Randwick NSW 2052, AUSTRALIA.

## Supplemental Figures

**Figure S1.**
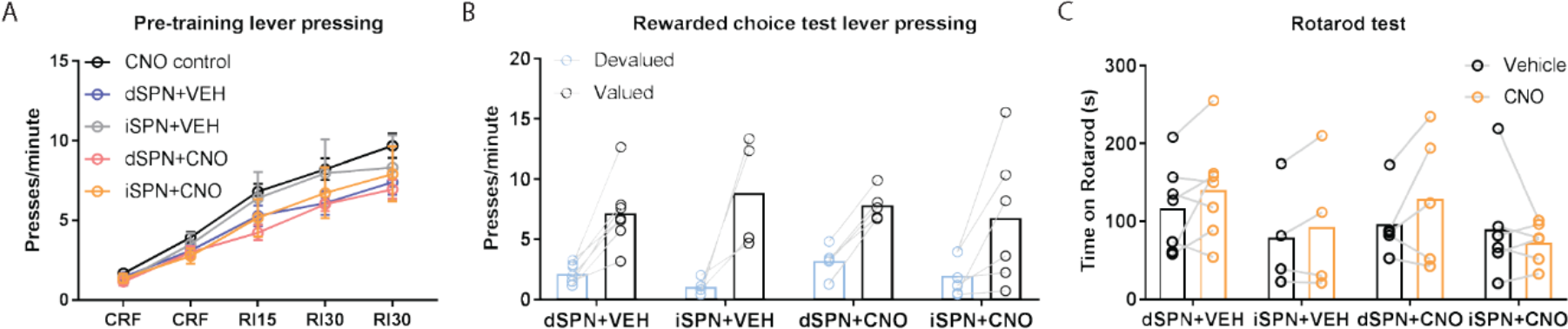
Bilateral chemogenetic inhibition of SPNs does not impair rewarded choice or rotarod performance. **A,** Mean (± SEM) lever presses per minute averaged across each day of instrumental pre-training for each group. **B,** Mean (± SEM) lever presses per minute on the devalued and valued lever for each rat in each group, averaged across two days of rewarded choice tests. **C,** Mean (± SEM) time spent on the rotarod for each rat in each group following either vehicle or CNO injections.

**Figure S2.**
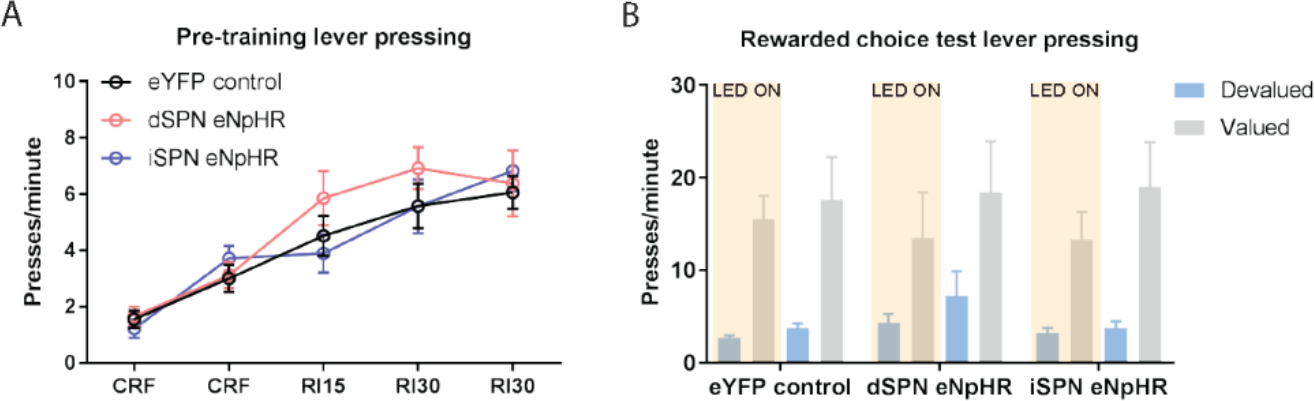
Optogenetic inhibition of SPNs does not impair rewarded choice. **A,** Mean (± SEM) lever presses per minute averaged across each day of instrumental pre-training for each group. **B,** Mean (± SEM) lever presses per minute on the devalued and valued lever for each group during LED ON (orange shaded) and LED OFF (non-shaded) periods, averaged across two days of rewarded choice tests.

